# Coalescent theory of the *ψ* directionality index

**DOI:** 10.1101/2025.06.18.660412

**Authors:** Egor Lappo, Noah A. Rosenberg

## Abstract

The *ψ* directionality index was introduced by Peter & Slatkin (*Evolution* 67: 3274-3289, 2013) to infer the direction of range expansions from single-nucleotide polymorphism variation. Computed from the joint site frequency spectrum for two populations, *ψ* uses shared genetic variants to measure the difference in the amount of genetic drift experienced by the populations, associating excess drift with greater distance from the origin of the range expansion. Although *ψ* has been successfully applied in natural populations, its statistical properties have not been well understood. In this paper, we define Ψ as a random variable originating from a coalescent process in a two-population demography. For samples consisting of a pair of diploid genomes, one from each of two populations, we derive expressions for moments 𝔼 [Ψ^*k*^] for standard parameterizations of bottlenecks during a founder event. For the expectation 𝔼[Ψ], we identify parameter combinations that represent distinct demographic scenarios yet yield the same value of 𝔼[Ψ]. We also show that the variance 𝕍[Ψ] increases with the time since the bottleneck and bottleneck severity, but does not depend on the size of the ancestral population; the ancestral population size affects *ψ* computed from many biallelic loci only through its contribution to the total number of loci available for the computation. Finally, we analyze the values of 𝔼[Ψ] computed from existing demographic models of *Drosophila melanogaster* and compare them with empirically computed *ψ*. Our work builds the foundation for theoretical treatments of the *ψ* index and can help in evaluating its behavior in empirical applications.

**Summary:** The statistic known as the “directionality index” examines variants shared between two populations with a goal of identifying which population has experienced a greater amount of genetic drift. This study develops theoretical predictions for the directionality index in coalescent models of pairs of populations descended from a common ancestral population. It determines the influence of bottlenecks, population growth, and population sizes on the directionality index. In *Drosophila melanogaster*, patterns in genomic data accord with the direction of the model predictions, with *ψ* suggesting a higher level of drift in European than in African populations.

## 1 Introduction

Inference of the demographic history of populations—including their population-size changes and relationships with other populations—is a major objective of statistical population genetics (e.g. Marchi et al., 2021). The combination of statistical methods based on coalescent theory with extensive genetic data has enabled researchers to investigate diverse features of demographic histories (e.g. Pool et al., 2010).

One of the most fundamental ways in which genetic data can be summarized for statistical analysis is by the site frequency spectrum (SFS), which counts the numbers of sites—typically single-nucleotide polymorphisms, or SNPs—that are present in different multiplicities in a sample (e.g. Wakeley & Hey, 1997; Achaz, 2009). Comparisons of the empirical SFS in a population to predictions of a coalescent model can detect phenomena such as bottlenecks, expansions, or selective sweeps (e.g. Ferretti et al., 2010; Ronen et al., 2013). The SFS has received extensive theoretical treatment under many demographic scenarios and has often been applied for inference in real populations (e.g. Nielsen et al., 2005; Thornton & Andolfatto, 2006).

In data from *multiple* populations, a joint SFS can be defined that records SNP allele frequencies in each population (Gutenkunst et al., 2009). A joint SFS enables inference of processes such as admixture, migration, and differences in selection between populations (Caicedo et al., 2007; Nielsen et al., 2009; Excoffier et al., 2013; Zhan et al., 2014; Arguello et al., 2019; Liu & Fu, 2020). In the setting of population pairs, Peter and Slatkin (2013) proposed a statistic, the *ψ* directionality index, which is computed from the joint SFS for the two populations. This index was designed for characterizing the process of range expansion, in which a population sequentially settles locations increasingly distant from its origin (e.g. Ramachandran et al., 2005; Excoffier & Ray, 2008; Excoffier et al., 2009).

In a range expansion, the leading edge of the expansion experiences stronger genetic drift relative to the point of origin (e.g. Hallatschek & Nelson, 2008; Slatkin & Excoffier, 2012; Peter & Slatkin, 2015; Peischl & Excoffier, 2016). In the genetic history of individuals at the edge of the expansion, the range expansion process can manifest as a sequence of population size bottlenecks, as increasingly distant geographic locations are settled (e.g. Deshpande et al., 2009; DeGiorgio et al., 2009, 2011).

For two populations that are part of the expansion, the *ψ* directionality index seeks to identify the direction of the expansion. The approach relies on the fact that if a given derived allele is shared between the two populations, then its frequency is expected to be higher in the derived population at the edge of the range expansion than in the source population (e.g. Edmonds et al., 2004; Klopfstein et al., 2006; Excoffier & Ray, 2008; Schlichta et al., 2022). Alleles at low frequency in the source population are likely to be lost during the expansion and therefore would not be shared. The derived population has a smaller founding population size than the source population, so that alleles—if they are not entirely absent—tend to possess greater frequencies. The *ψ* index considers the population differences of allele frequencies specifically in the shared genetic variation between the two populations.

Among pairwise quantities that can be computed as summary statistics useful for interpreting population-genetic data (e.g. *F*_*ST*_), the *ψ* index stands out as a signed quantity. For two populations *A* and *B*, the order of the populations matters, with *ψ*(*A, B*) = −*ψ*(*B, A*). Therefore, whereas *F*_*ST*_ is often seen as a genetic measure of distance, *ψ* is akin to a vector directed from one population to another (see also Peter and Slatkin, 2013, Figs. 5, 7).

The *ψ* index was first defined by Peter and Slatkin (2013), who developed a method that integrates information about pairwise *ψ* with geographic distances between sampling locations to identify coordinates of the expansion origin. They then applied it to simulated scenarios including isolation-by-distance and range expansion on a grid of populations, as well as to complex configurations involving migration barriers.

Peter and Slatkin (2015) then studied theoretical properties of *ψ* in a discrete time-expansion model. The model consisted of a linearly arranged set of demes with equal population size, with a single deme *d*_0_ settled initially and the rest of the demes empty. At an integer timepoint *t*, a new deme *d*_*t*_ is settled by individuals from the previous deme *d*_*t−*1_. The quantity of interest was *ψ*(*d*_0_, *d*_*t*_) at time *t* between the origin deme and the leading edge of the expansion. Peter and Slatkin (2015) showed that in the model, the expected value of *ψ* between the source and the leading edge of the expansion—which has experienced a sequence of founder events—depends on the relative founder sizes of settlement events (the fraction of individuals selected from deme *d*_*t−*1_ to settle *d*_*t*_) and the number of founder events, equal to *t* in their scaling of time. Peter and Slatkin (2015) used *ψ* to identify the expansion origin for natural populations of *Arabidopsis thaliana*, which they presumed to have expanded spatially in a manner compatible with a linear arrangement of demes.

Several recent uses of *ψ* have since sought to examine scenarios where, instead of an expansion over a linear spatial dimension, an expansion involves pairwise computations for a small number of discrete demes, as few as 2. For example, Zhan et al. (2014) examined the expansion of monarch butterflies from North America to South America, the Pacific, and Europe, computing *ψ* between a source population in North America and a destination population elsewhere. Puckett and Munshi-South (2019) examined the expansion of brown rats from Eastern Asia to the Middle East, the Middle East to Europe, and Europe to North America, computing *ψ* between pairs of populations in two different geographic regions. Ioannidis et al. (2021) similarly used pairwise values of *ψ* between pairs of human populations of different Pacific islands to understand sequences of events in the human settlement of the region.

In this paper, building from the interest in using *ψ* for expansions involving small numbers of discrete populations rather than many demes along a spatial continuum, we examine the *ψ* statistic theoretically in the simplest discrete–deme structured population: a pair of populations. We define Ψ as a random variable arising from the coalescent process (Section 2) and derive expressions for moments of ΨΨ under the coalescent. We focus on the scenario in which a single diploid individual is sampled in each a pair of populations. In Section 3, we consider specific commonly used parameterizations of range expansions in the setting of population pairs, explicitly incorporating exponential growth, bottlenecks, and instantaneous bottlenecks (Figure 1). In Section 4, we explore theoretical predictions for the expectation 𝔼[Ψ] and variance 𝕍[Ψ], interpreting them in terms of the reliability of inferences and the identifiability of demographic scenarios. In Section 5, we use the central limit theorem to analyze the sample variance of the *ψ* index computed from many SNPs. Finally, in Section 6, we show how our results can be used in the evaluation of empirical inferences of demographic parameters for real populations.

**Figure 1:**
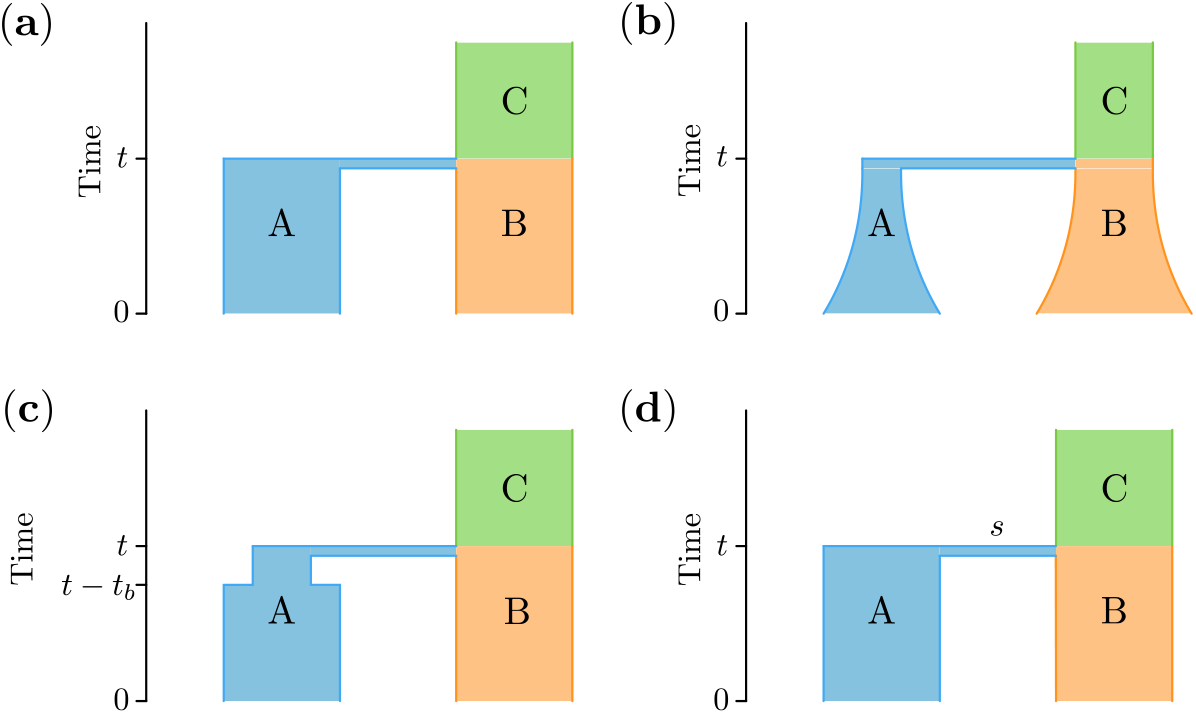
Four demographic scenarios. (a) Population split: an ancestral population C splits into two populations A and B at time *t*. (b) Exponential growth: after the split, populations A and B grow at rates *r*_*A*_ and *r*_*B*_, respectively. (c) Bottleneck: after the split, population A goes through a bottleneck of population size *N*_*b*_ and duration *t*_*b*_. (d) Instantaneous bottleneck: after the split, population A goes through a burst of coalescences of strength *s*. Times *t* and *t− t*_b_ are measured in generations back from the present.

## 2 Coalescent-based definition of the *ψ* index

### 2.1 *ψ* for a pair of genomes

The directionality index *ψ* is a two-population statistic computed from allele frequencies for a set of biallelic SNPs. For the rest of the paper, we assume that the derived and ancestral alleles are known for each SNP, and we call the SNP *shared* between two populations if the derived allele is present in at least one copy in both populations and the SNP is polymorphic in the pooled pair of populations.

Suppose now that we know allele frequencies for a set of SNPs in two populations A and B. In its most general form, the value of the *ψ* index is then defined as

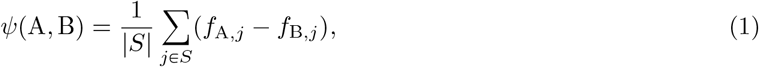

where *S* is the set of SNPs *shared* between the two populations, *f*_A,*j*_ is the frequency of the derived allele of SNP *j* in population A, and *f*_B,*j*_ is its frequency in population B (Peter & Slatkin, 2015, eq. 1).

We proceed by focusing on the simplest case in which *ψ* can be meaningfully studied in two populations. In particular, if allele frequencies are computed using a single diploid individual sampled from each population, and if shared SNPs are identified based on this pair of individuals, then the expression for *ψ* reduces to

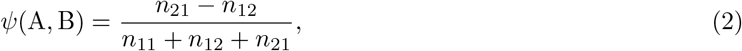

where *n*_*ij*_ is the number of SNPs that have *i* copies of the derived allele in the individual from population A and *j* copies in the individual from population B, and *i* and *j* can each equal 0, 1, or 2.

### 2.2 The random variable Ψ under the coalescent

We now analyze the *ψ* index as a random variable under the coalescent. We use the notation Ψ to distinguish the theoretical random variable for the directionality index from the empirical *ψ* computed from data.

We assume that all SNPs are unlinked, so that coalescent trees for different SNPs are independent. We also assume that SNPs obey the standard infinitely-many-sites mutation model (Durrett, 2008, p. 29), such that each SNP results from a single mutation on a coalescent tree. Finally, we assume that we have specified a demographic history for populations A and B (Figure 1), and that a single diploid individual is sampled from each population. Such a sample configuration—one diploid individual with sample size 2 alleles in each population—allows us to use the simplified eq. (2).

Conditional on the demography and a sample of size 2 from each of a pair of populations, the coalescent model defines a probability distribution over the genealogies of lineages from A and B. In this framework, we can determine the expectation 𝔼[Ψ](*A, B*) of the directionality index under the coalescent model for a single SNP shared between populations A and B. We use 𝔼 [Ψ^*k*^] with *k* > 1 to denote higher moments of the random variable Ψ under the coalescent.

To compute 𝔼[Ψ], we consider probabilities under the coalescent model of entries in the joint site frequency spectrum for populations A and B, conditional on the demography:

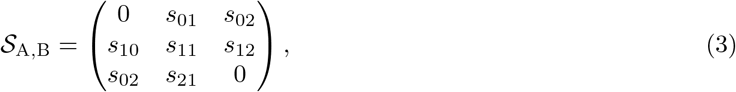

where *s*_*ij*_ is the probability that a randomly sampled mutation—that is, the derived variant of a random SNP on the genealogy of four lineages—occurs in *i* copies in population A and in *j* copies in population B. For example, *s*_12_ is the probability that for a random SNP, the diploid sample from population A has one ancestral and one derived allele, and the sample from population B is homozygous with two copies of the derived allele. As a shorthand, we will say that such a SNP has “type 12,” and we indicate other elements of the site frequency spectrum similarly.

Suppose now that we have sampled a single *shared* SNP. First, the probabilities of a SNP having a specific type are obtained from the site frequency spectrum *𝒮* by conditioning on being shared,

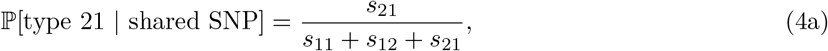

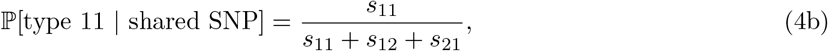

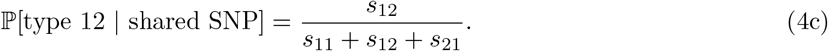

Further, taking the total number of sampled SNPs to be 1 in eq. (2), we know that the value of Ψ is constant for all SNPs of the same type, with

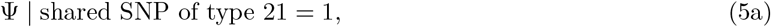

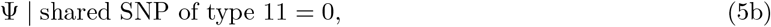

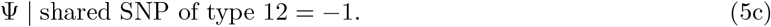

Combining eq. (4) with eq. (5), we can write the definition of random variable Ψ for a single shared SNP:

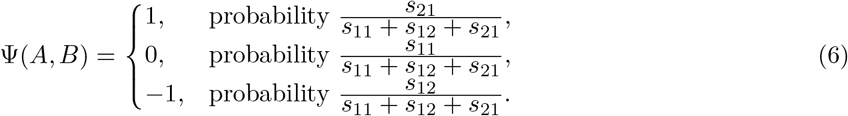

The expectation of Ψ can then be straightforwardly computed as

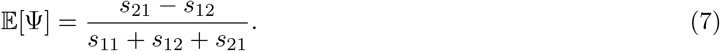

The second moment of Ψ is

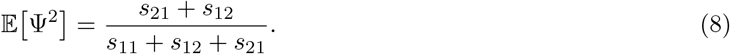

The higher moments of *ψ* can be computed similarly, with

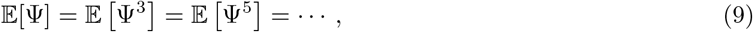

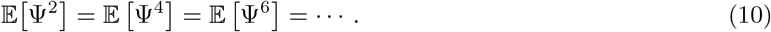

In the remainder of this paper, we discuss only the expectation 𝔼[Ψ] and the variance 𝕍[Ψ] = 𝔼 [Ψ^2^] − 𝔼[Ψ]^2^, as the other moments can be obtained from these cases.

The only remaining quantities we need are the *s*_*ij*_: the probabilities that a randomly chosen SNP has *i* copies of the derived allele in the sample of size 2 from population A and *j* copies in the sample of size 2 from population B, with (*i, j*) = (1, 1), (1, 2), or (2, 1). In other words, under a coalescent-based demographic model with the infinitely-many-sites mutation, we seek to compute, as a fraction of all SNPs, the number that occur on genealogical branches ancestral to *i* copies in population A and *j* copies in population B.

In a random genealogy, the expected total number of SNPs with type *ij* is Θ𝔼[*L*_*ij*_]/2, where *L*_*ij*_ is the total length of branches ancestral to *i* lineages from population A and *j* lineages from population B, and Θ/2 is the Poisson mutation rate along a branch. 𝔼[*L*_*ij*_] is computed by considering each topology separately:

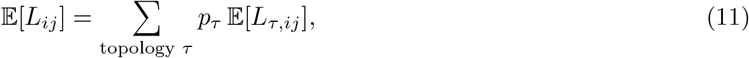

where *p*_*τ*_ is the probability that topology *τ* occurs and *L*_*τ,ij*_ is the length of branches ancestral to *i* lineages from A and *j* lineages from B in genealogies with topology *τ*. The value of *s*_*ij*_ is then proportional to

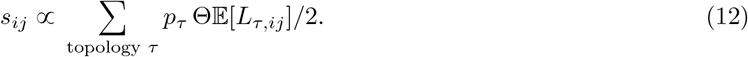

Because we regard lineages within populations as exchangeable—so that we do not distinguish between two lineages from the same population—six topologies must be considered in eq. (12) (Figure 2). We denote the six topologies *α, β, γ, δ, ε, ζ*. The topology probabilities *p*_*τ*_ and the expected branch lengths 𝔼[*L*_*τ,ij*_] can be computed for various demographic models, so that eq. (12) can be calculated and hence also eq. (7) and eq. (8). In the next section, we compute these quantities for simple models representing a founder effect.

**Figure 2:**
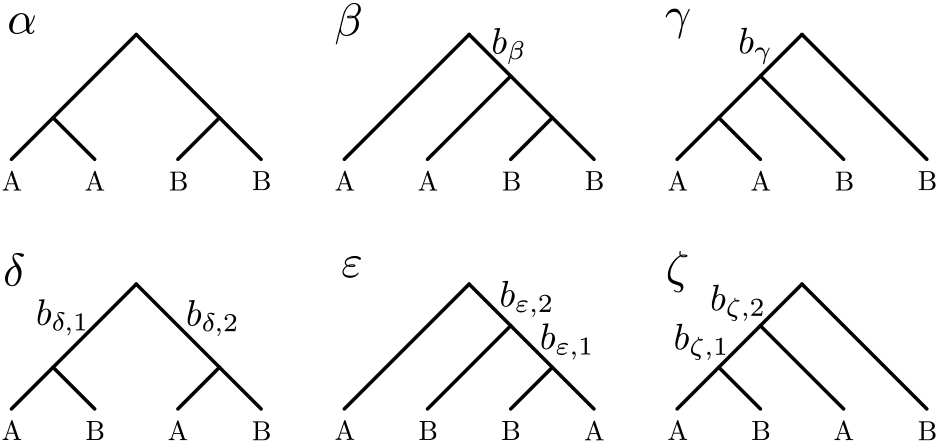
The six tree topologies possible for samples of two lineages each from two populations A and B. Trees are labeled by Greek letters. Branches relevant to the calculation of 𝔼[Ψ]—branches that are ancestral to lineages from both populations—are labeled by *b*_*β*_, *b*_*γ*_, etc.

## 3 Expectation and variance of Ψ for specific demographic models

The exact expressions for topology probabilities and branch lengths in eq. (12) depend on specific parameterizations of the demographic history. In this section we derive expressions for *s*_*ij*_ and moments of Ψ in eqs. (7) and (8) for the four demographies shown in Figure 1.

### 3.1 Population split

We first consider a simple population split demography (Figure 1a). A single ancestral population C of size *N*_C_ splits into two populations *t* generations ago. The two resulting populations A and B have sizes *N*_A_ and *N*_B_ individuals, respectively.

To compute 𝔼[Ψ] and 𝕍[Ψ], we compute topology probabilities and relevant branch length expectations for each topology in Figure 2. Computations with topology *α* are not needed because this topology cannot generate shared polymorphisms. For topology probabilities *p*_*β*_ and *p*_*γ*_, we must further distinguish the tree topologies based on the population in which the “cherry coalescence” happens. We denote by *p*_*β*,B_ the probability of the topology *β* in which the coalescence (B,B) happens in population B; if the coalescence (B,B) happens in the ancestral population C, then we denote the probability by *p*_*β*,C_, with similar notation for topology *γ*. These additional topology labels are depicted in Figure 3.

**Figure 3:**
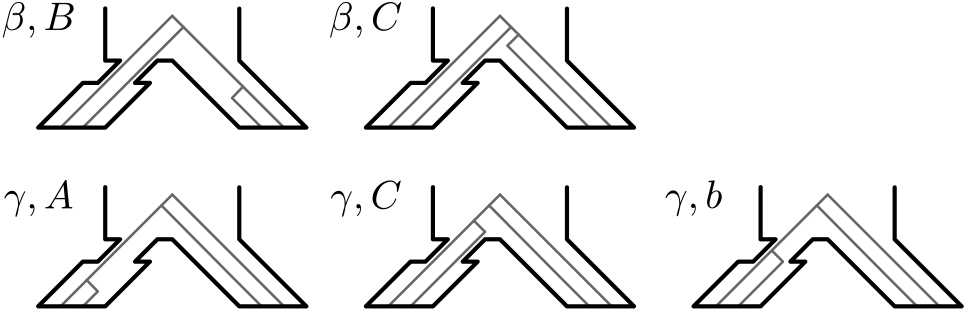
Distinguishing locations of the cherry coalescence for topologies *β* and *γ*. The ancestral population is C and the descendant populations are A (left) and B (right).

First, we compute *p*_*β*,B_. The two lineages from population B must coalesce in population B; the probability of this event is 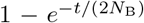. The two lineages from population A must not coalesce until they enter population C; the probability of that event is 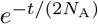. As a result, two lineages from A and one lineage from B enter population C. In population C, lineages from A and B must coalesce first; the probability of this event is 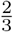; in the remaining 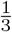 of cases, the two A lineages coalesce first. As a result, we obtain 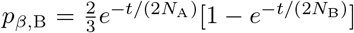. This derivation modifies the two-population calculation of Tajima (1983) by allowing for different population sizes for populations A and B rather than assuming their exchangeability. Using the same logic for other tree topologies, we obtain the following probabilities:

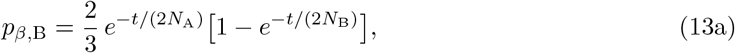

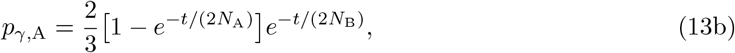

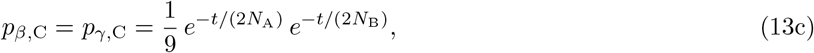

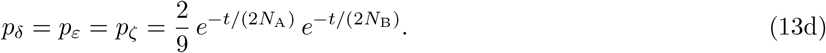

For *β* and *γ*, summing the probabilities of the two cases, we get

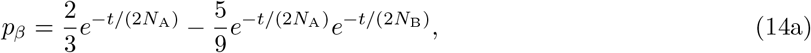

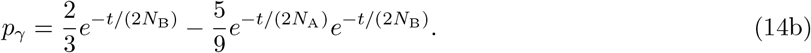

We now compute expected lengths of relevant branches for specific sample configurations. All branches below the gene tree root that are shared by at least one pair of lineages from different populations are labeled in Figure 2. For example, branch *b*_*β*_ is ancestral to two lineages from population B and one lineage from population A for topology *β*; similarly, branch *b*_*ζ*,1_ is ancestral to one lineage from population A and one lineage from population B if the genealogy has topology *ζ*.

With eq. (12), we obtain equations for entries of the expected joint site frequency spectrum 𝒮_A,B_:

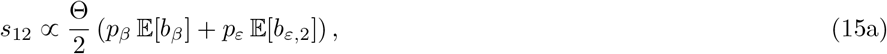

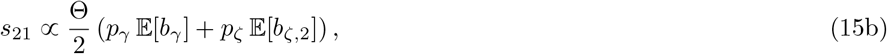

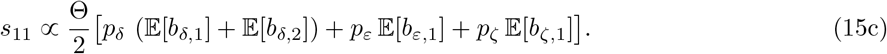

In the final expressions for moments of Ψ (eqs. (7) and (8)), the mutation rate cancels because all branches that can generate shared sites can only appear in population C, so that Θ = 4*N*_C_*µ* in all parts of eq. (15).

We are now left with calculating the expected branch lengths. Because polymorphisms shared between populations A and B can only result from mutations in the ancestral population C, our branch length calculations need only consider coalescent theory in a single population of size *N*_C_ individuals. In particular, expectations of branch lengths *b*_*β*_ and *b*_*γ*_ are equal to the expectation of the time 𝔼[*T*_2_] to coalescence of two lineages in the diploid population of size *N*_C_, so that

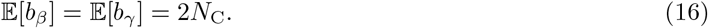

Similar logic applies to trees *ε* and *ζ*, with expectations of *b*_*ε*,1_ and *b*_*ζ*,1_ equaling the expectation of the time *T*_3_ to the first coalescence with three lineages,

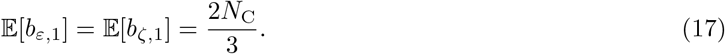

The lengths of *b*_*ε*,2_ and *b*_*ζ*,2_ are again proportional to 𝔼[*T*_2_] as in eq. (16),

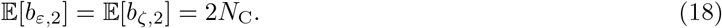

Finally, the expected length of branches *b*_*δ*,1_ and *b*_*δ*,2_ together is equal to 2𝔼[*T*_2_] + 𝔼[*T*_3_],

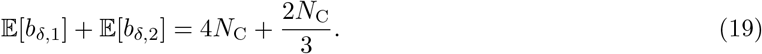

Finally, we can substitute expressions for the branch lengths (eqs. (16) to (19)) and the topology probabilities (eq. (13)) into eq. (15) to obtain expressions for SFS entries *s*_*ij*_. We then plug these quantities into eqs. (7) and (8) to obtain

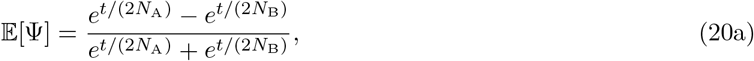

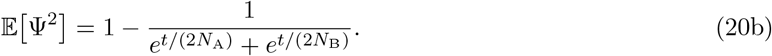

The variance of Ψ is then

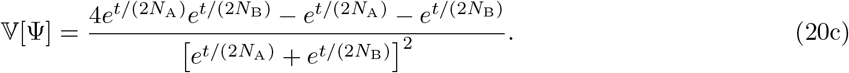

Examining the expressions in eqs. (20a) and (20c), we see that both the mean of Ψ and the variance of Ψ do not depend on the size *N*_*C*_ of the ancestral population. We also observe that if the population sizes are equal, *N*_A_ = *N*_B_, then 𝔼[Ψ] = 0. Moreover, examination of eq. (20) can directly bound the expectation and variance. As 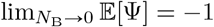 and 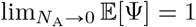, we have -1 < 𝔼[Ψ] < 1; for variance, we have 0 < 𝕍[Ψ] < 1 because, on one hand, lim_*t*_𝕍[Ψ] = 0, and on the other hand,

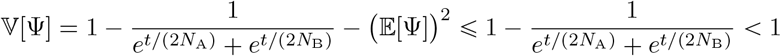

for *N*_A_, *N*_B_ > 0 and *t* ⩾ 0. We will use these facts in our exploration of the behavior of Ψ in Section 4.

Informally, we can write eq. (20a) as

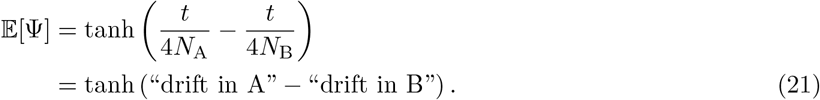

In this simple model, the “amount of drift” is that of a neutral population of size *N*_A_ (or *N*_B_) evolving for *t* generations. However, by treating *N*_A_ and *N*_B_ as effective population sizes, a variety of demographic scenarios that include population growth or bottlenecks can be considered. In subsequent subsections, we explicitly parameterize models with population size changes and present modified versions of eq. (20).

### 3.2 Exponential growth

We next consider populations A and B evolving under the classic exponential growth model. A and B begin exponential growth immediately after splitting from the ancestral population C, as shown in Figure 1b.

Let population A have size *N*_A,0_ at the present time, such that its population size over time is

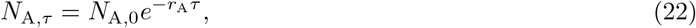

where *τ* is time, measured in generations from the present into the past, and *r*_A_ is the growth rate. Eq. (22) is defined such that if *r*_A_ > 0, then population A is increasing in size forward in time. If population A has size *N*_A,*t*_ immediately after the split, then the growth rate can be computed from eq. (22) as

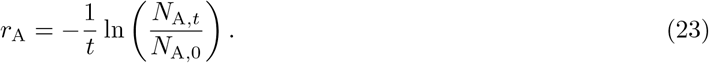

Slatkin and Hudson (1991) showed that for a pair of lineages, the coalescent in a growing population of size *N*_A,*τ*_ is equivalent to the coalescent in the constant population of size *N*_A,0_, with time rescaled by

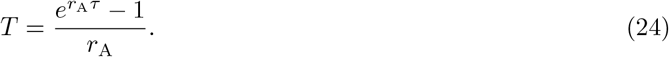

Hence, the probability that two lineages coalesce in the first *t* generations in the population of size *N*_A,*τ*_ is

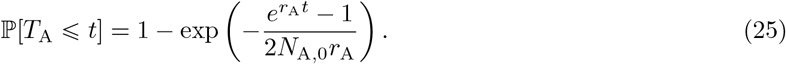

A corresponding equation holds for population B.

We can repeat the calculations of tree topology probabilities in eqs. (13) and (14) by replacing the constant-size coalescence probability 1 − exp[−*t*/(2*N*)] by the quantity in eq. (25). As a result, we obtain the following expressions for expectation and variance of *ψ* under the exponential growth model:

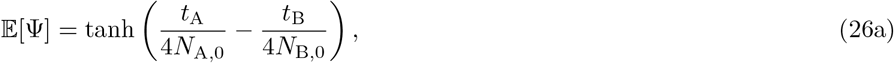

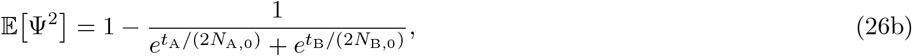

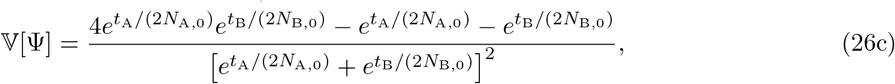

where we have introduced a shorthand notation

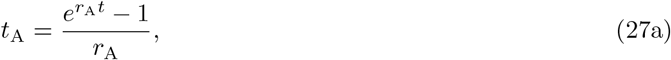

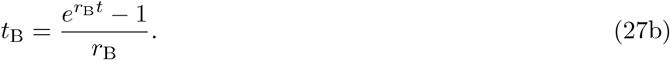

If only one population is subject to exponential growth, then expressions for 𝔼[Ψ] and 𝕍[Ψ] can be found by taking the limits in eq. (26) as the growth rate approaches zero. For example, if *r*_B_ = 0, then we have

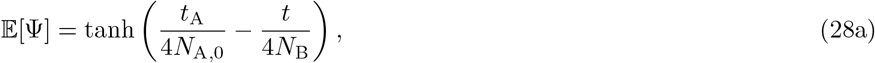

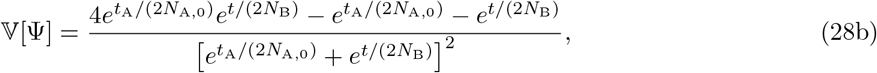

where again *t*_A_ is defined by eq. (27a).

### 3.3 Bottleneck

For our third model, we assume that immediately after the split, population A goes through a bottleneck of length *t*_b_ with constant population size *N*_b_, as shown in Figure 1c. This type of model has been used in studies of human expansion from Africa (DeGiorgio et al., 2009, 2011).

The calculations of 𝔼[Ψ] and 𝕍[Ψ] are similar to those in Section 3.1, except that the topology probabilities differ: we consider special cases for topologies *β* and *γ* (Figure 2). For *β*, we distinguish between coalescent genealogies in which the node (B,B) is located in population B (*p*_*β*,B_) and in population C (*p*_*β*,C_). For *γ*, we distinguish between genealogies in which the node (A,A) occurs in population A after the bottleneck (*p*_*γ*,A_), during the bottleneck (*p*_*γ*,bot_), and in population C (*p*_*γ*,C_). The probabilities are:

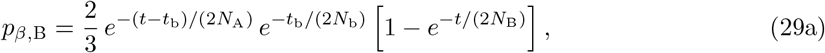

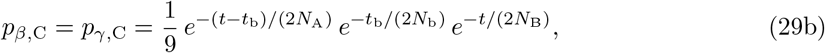

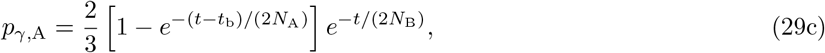

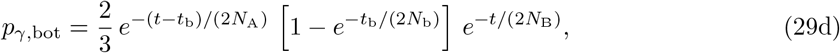

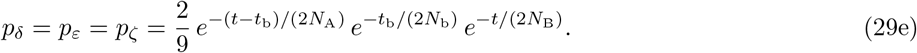

The expressions for the moments of Ψ are

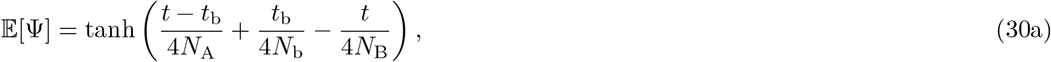

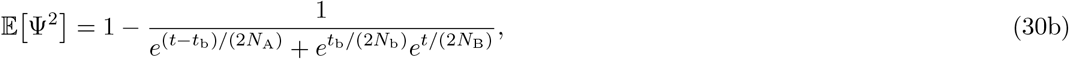

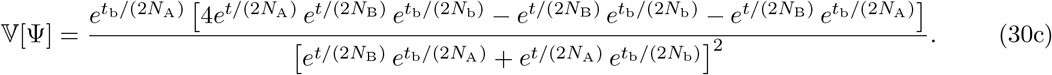

If *N*_b_ = *N*_A_, then eq. (30) reduces to eq. (20). If *t*_b_ = 0, then expressions in eq. (30) match eq. (20) irrespective of the value of *N*_b_.

### 3.4 Founder effect

The final model that we consider is a model that has been proposed for simplifying the modeling of founder effects. Instead of a prolonged bottleneck, we introduce an *instantaneous bottleneck* into population A (Figure 1d). An instantaneous bottleneck is defined as a burst of coalescences; mathematically, two lineages going through an instantaneous bottleneck of strength *s* behave as if going through *s* (imaginary) generations of drift in the population of final size *N*_A_. Instantaneous bottlenecks are typically used in situations where the bottleneck is short enough such that the possibility of mutations happening *during* the bottleneck can be disregarded (Galtier et al., 2000; Bunnefeld et al., 2015). In practice, this scenario could correspond to a low number of lineages from population C settling the whole population A that exists after the split.

Similarly to Section 3.3, we adjust the tree topology probabilities to reflect the demography in Figure 1d:

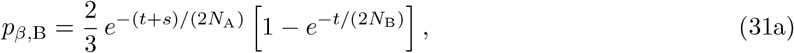

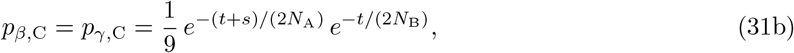

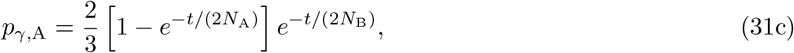

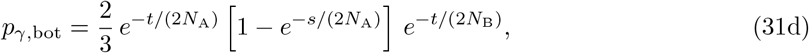

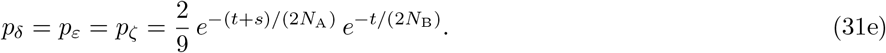

The expressions for the moments of Ψ in this case are:

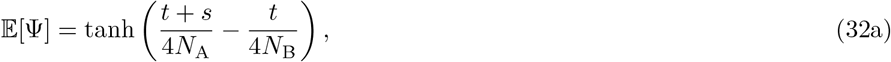

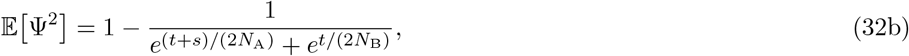

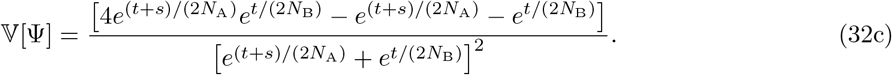

These expressions reflect the fact that the “strength” of the bottleneck depends on its duration *t*_b_ and population size *N*_b_ only through the ratio *t*_b_/(2*N*_b_), as captured by the parameter *s*. If *s* = 0, then eq. (32) reduces to eq. (20).

## 4 Illustrations of 𝔼[Ψ] and 𝕍[Ψ]

To illustrate our theoretical expressions, we plot 𝔼[Ψ] and 𝕍[Ψ] for a range of parameter values. For these plots, we use the instantaneous bottleneck formulation of Section 3.4, as it has only five parameters *t, s, N*_A_, *N*_B_, and *N*_C_ instead of six, as in Section 3.2 or Section 3.3.

Figure 4 shows 𝔼[Ψ] and 𝕍[Ψ] for varying *t* and *s* and fixed population sizes *N*_A_ = 400, *N*_B_ = 600, and *N*_C_ = 1000. The behavior of 𝔼[Ψ] confirms the intuition in eq. (21): as the bottleneck strength *s* increases, population A accumulates larger amounts of drift due to increased probability of coalescence in the bottleneck (eq. (32a)), leading to higher positive values of 𝔼[Ψ]. Similarly, increasing *t* leads to more drift in population A because *N*_A_ < *N*_B_, and A accumulates drift at a higher rate than B. Note that if we were to set *N*_A_ = *N*_B_ in eq. (32a), then the value of 𝔼[Ψ] would not depend on the time *t* since the bottleneck.

**Figure 4:**
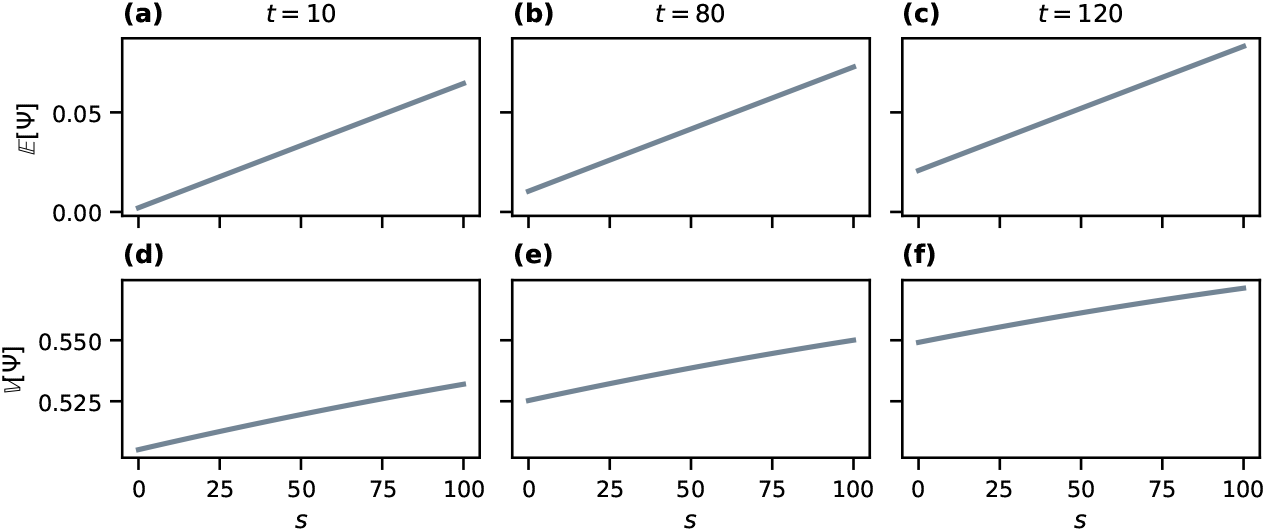
Values of 𝔼[Ψ] and 𝕍 [Ψ] under the founder effect demography of Figure 1d. The population sizes are constant, with *N*_A_ = 400, *N*_B_ = 600 and *N*_C_ = 1000. The values are computed using eq. (32a) for 𝔼[Ψ] and eq. (32c) for𝕍 [Ψ]. (a) 𝔼[Ψ] for varying *s, t* = 10. (b) 𝔼[Ψ] for varying *s, t* = 80. (c) 𝔼[Ψ] for varying *s, t* = 120. (d) 𝕍 [Ψ] for varying *s, t* = 10. (e) 𝕍[Ψ] for varying *s, t* = 80. (f) 𝕍 [Ψ] for varying *s, t* = 120.

The variance 𝕍[Ψ] also increases with both the time *t* since the bottleneck and the bottleneck strength *s*, with stronger dependence on *t* compared to 𝔼[Ψ]. As *s* or *t* increases, the probability of observing a SNP of type 11, *s*_11_, decreases, as a type 11 SNP requires all four lineages from A and B to persist into population C without coalescing, whereas type 12 and type 21 SNPs can be produced with only three lineages persisting into population C. SNPs of type 11 can appear *only* in genealogies *δ, ε*, and *ζ* where all coalescences happen in ancestral population C, and the probability of observing these genealogies is small for large *s* and *t*. The second moment 𝔼 [Ψ ^2^] = (*s*_21_ + *s*_12_)/(*s*_12_ + *s*_21_ + *s*_11_) then grows large, increasing the variance.

We can use our theoretical results to analyze identifiability of demographic scenarios with the *ψ* index. In analyses of genetic data, a positive *ψ* is used to claim that the population A is located further from the source of the range expansion (Peter & Slatkin, 2013), with the population with more drift experiencing bottlenecks during founder events. However, this logic does not account for the possibility that other demographic scenarios could generate an identical value of *ψ*. Figure 5 provides an example of this phenomenon by showing that if the population size *N*_B_ is sufficiently small relative to *N*_A_, then 𝔼[Ψ] could be negative even in the presence of a bottleneck in population A. Figure 6 shows the dependence of 𝔼[Ψ] on *N*_*B*_ and *t*. If *N*_B_ is small enough in relation to *N*_A_, then the value of 𝔼[Ψ] can decrease with increasing *t* and can even reverse its sign. The negative value of the directionality index then obscures the ancient bottleneck in A.

**Figure 5:**
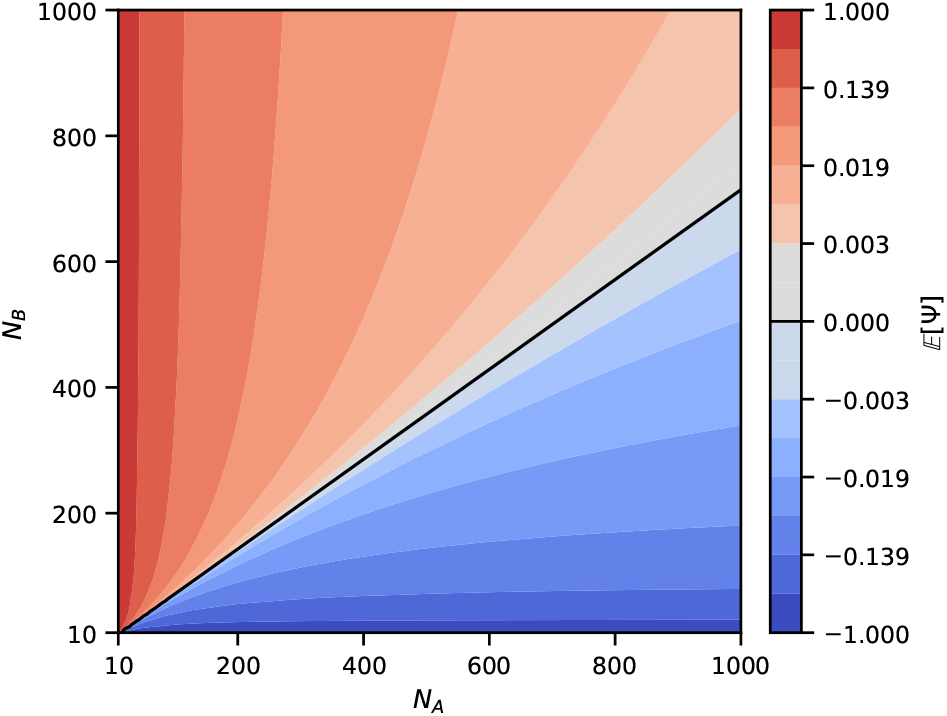
Theoretical values of 𝔼[Ψ] (eq. (32a)) for varying values of *N*_A_ and *N*_B_ in the founder effect demography of Figure 1d. Parameters *s* and *t* are fixed, with *s* = 20 and *t* = 50. The black line shows parameter sets (*N*_A_, *N*_B_) for which 𝔼[Ψ] = 0.

**Figure 6:**
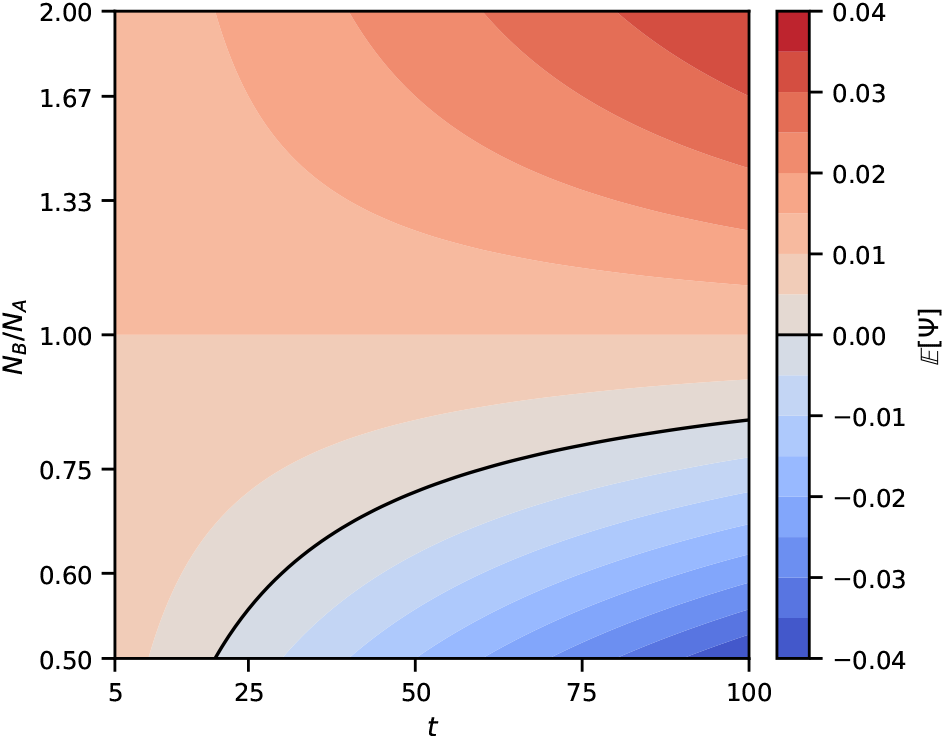
Theoretical values of 𝔼[Ψ] (eq. (32a)) for varying values of *t* and *N*_B_ in the founder effect demography of Figure 1d. Parameters *s* and *N*_A_ are fixed, with *s* = 20 and *N*_A_ = 500. The black line shows parameter sets (*t, N*_B_) for which 𝔼[Ψ] = 0.

## 5 Sampling theory of Ψ

We have demonstrated that the expectation 𝔼[Ψ] and variance 𝕍[Ψ] do not depend on the ancestral population size *N*_C_. In this section, we show that our confidence in the value of *ψ* computed from SNP data does depend on *N*_C_ through sample variance.

The random variable Ψ and its associated quantities 𝔼[Ψ] and 𝕍[Ψ] refer to the directionality index for a single SNP under the coalescent. In a data analysis, *ψ* is computed using many shared SNPs across the genome, say *n*. Denote by 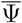 the (random) many-SNP *ψ* index, signifying that this quantity can be seen as the mean of many single-SNP observations of *ψ*. For *n* sampled independent shared SNPs, the central limit theorem states that the resulting 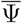 approaches a Gaussian distribution with variance proportional to 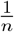,

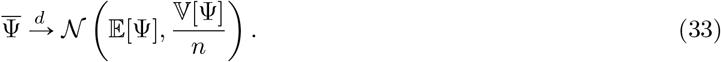

As the number of sampled shared SNPs increases, the probability that 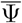 is close to its mathematical expectation 𝔼[Ψ] increases.

Under the infinitely-many-sites model, the number of shared SNPs *n* is itself a random variable that depends on mutation rate *µ* and the ancestral population size *N*_C_, as shared SNPs reflect mutations in the ancestral population. More precisely, 𝔼[*n*] = Θ𝔼[*L*]/2, where Θ = 4*N*_C_*µ* is the scaled mutation rate in a diploid population of *N*_C_ individuals, and 𝔼[*L*] is the expected length of branches that can yield shared SNPs, in units of 2*N*_C_ generations. All branches that can generate shared SNPs were identified in Figure 2, so we can use eqs. (16) to (19) to write an equation for 𝔼[*L*]:

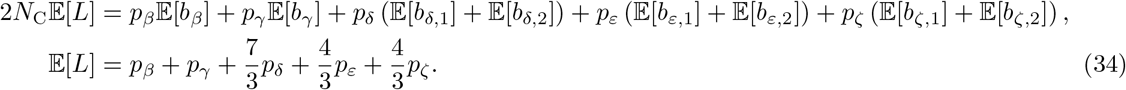

For example, with the founder effect model of Section 3.4, we get

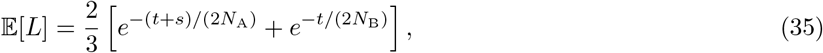

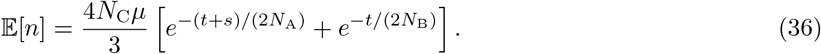

The expected number of shared SNPs depends linearly on the ancestral population size, and *N*_C_ then affects the number of shared SNPs available for empirical analyses using the directionality index.

Eq. (36) specifies the dependence of the random variable *n* on the demographic parameters. For example, we can see that stronger bottlenecks—higher values of *s*—lead to an increase not just in the variance 𝕍[Ψ] (Figure 4), but also in the sample variance by decreasing the number of available shared SNPs, 𝔼[*n*].

## 6 Application to Out-of-Africa expansion of *D. melanogaster*

To test our coalescent-based predictions for *ψ*, we compute *ψ* for a specific demographic event in two ways. First, we use eq. (1) to compute *ψ* directly from genotypes in natural populations. Second, we make use of existing estimates of demographic parameters to evaluate our equations for 𝔼[Ψ] and 𝕍[Ψ].

The demographic scenario we consider here is the Out-of-Africa expansion of *Drosophila melanogaster*. It is generally agreed that the modern European populations trace to a small founding population as the species range expanded from Africa (e.g. Stephan & Li, 2007; Arguello et al., 2019). As the founder event was directed from Africa to Europe, we expect to see *ψ*(Europe, Africa) > 0.

### 6.1 *ψ* computed from sequence data

For our empirical computations, we evaluated *ψ* from the sequences of intronic and intergenic X-chromosomal loci used for demographic inference by Li and Stephan (2006), Laurent et al. (2011), and Duchen et al. (2013), originally obtained by Glinka et al. (2003) and Ometto et al. (2005). We downloaded sequences of X-chromosomal loci from the European Nucleotide Archive (ebi.ac.uk/ena), sequence IDs AJ568984 to AJ571588 and AJ568984 to AJ571588 (originally deposited by Glinka et al. (2003) and Ometto et al. (2005)).

Each locus had nucleotide data in the form of a single (haploid) genotype for a set of inbred lines from European (Netherlands, NTH) and African (Zimbabwe, ZW) populations of *D. melanogaster*, as well as for a single line from a North American population of *D. simulans*. The genetic sequence for each line was haploid due to the sequencing being performed with homozygous inbred lines. The total number of X-chromosomal loci was 229, with locus sequence lengths ranging from 210 to 784 nucleotides (median 563).

The *D. simulans* sequence was used in place of the ancestral genotype in the analysis. Across the 229 loci, the maximum number of lines sequenced for the Netherlands population was 12 and the minimum number of lines sequenced was 10. For the Zimbabwe population, the maximum was 12 and the minimum was 9.

Separately for each locus, we used *MUSCLE v5*.*1* (Edgar, 2004) with default settings (`-perturb 0 -perm none -consiters 2 -refineiters 100`) to perform a joint multiple sequence alignment for the lines from the NTH and ZW populations of *D. melanogaster* as well as the *D. simulans* line.

To compute *ψ* for a set of loci, we generated a sample of 1000 sets of four lines, two from the NTH population and two from the ZW population. The sets of four were sampled with replacement, but each set had two distinct NTH lines and two distinct ZW lines (“distinct” here refers to distinct sample labels, not to distinctness of the genotypes).

For each set of four lines together with the *D. simulans* line, we discarded sites that had insertions or deletions in the alignment of five sequences. We next discarded invariable sites as well as sites with three or more distinct alleles. Next, we discarded sites that failed to meet a sharing criterion. In particular, we kept only those shared (biallelic) sites in the sense of our definition in Section 2, requiring the derived allele to be present in at least one copy in both NTH and ZW populations and to be polymorphic in the pooled pair of populations. We then computed *ψ* using eq. (2) by sampling a single site from the final set of shared sites.

To understand the uncertainty in the *ψ* computation that arises from differences in evolutionary history across loci, we analyzed 1000 bootstrap replicate datasets, where each bootstrap replicate involves a resample of 229 loci. In particular, in each bootstrap replicate, we first sampled 229 loci with replacement. Next, we generated 1000 sets of four lines and computed *ψ*, as described in the previous two paragraphs.

For each bootstrap replicate, we averaged *ψ* over the 1000 values, each obtained from a random set of four lineages, obtaining a mean *ψ* for that replicate. We also obtained a variance across the 1000 values.

Considering the 1000 bootstrap replicates, the median of the mean *ψ* values was

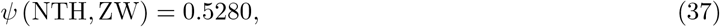

with 95% of mean *ψ* values lying in the interval (0.4860, 0.5710).

The median across 1000 bootstrap replicates of the variance of *ψ* was equal to

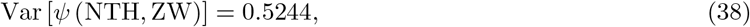

and the interval containing the variance values from 95% of the replicates was (0.4811, 0.5666).

### 6.2 𝔼[Ψ] computed from demographic estimates

We now compare empirical *ψ* values with the *ψ* values predicted by demographic models; we use demographic models that have been inferred in studies of the European founder event in *D. melanogaster*.

Multiple studies have estimated population sizes and divergence times for *D. melanogaster*. In particular, Li and Stephan (2006) used a maximum likelihood method based on the joint SFS, and Laurent et al. (2011) and Duchen et al. (2013) used approximate Bayesian computation. All three studies used the same set of X-chromosomal sequences from Glinka et al. (2003), with the Netherlands representing Europe and Zimbabwe representing Africa.

The study of Laurent et al. (2011) incorporates Asian samples, and the study of Duchen et al. (2013) adds North American samples, but here we focus on subsets of the inferred demographic parameters in these studies, specifically on the divergence of African and European populations shared by all three studies.

The modes of parameter estimates from the three papers are summarized in Table 1. The model of Li and Stephan (2006) assumes a prolonged bottleneck of constant size in the European population. Laurent et al. (2011) and Duchen et al. (2013) instead assume exponential growth in Europe. All three models assume constant population size in the African population after the split with the European population.

**Table 1:** Inferred demographic parameters reported for European and African *Drosophila melanogaster* populations in previous studies that used X-chromosomal loci, and corresponding model-predicted values of 𝔼[Ψ] and 𝕍 [Ψ]. In all studies, ten generations per year are assumed; we report times in generations.

| Paper | $t$ | $N_{\text{AFR}}$ | European bottleneck | $\mathbb{E}[\Psi](\text{EUR}, \text{AFR})$ | $\mathbb{V}[\Psi](\text{EUR}, \text{AFR})$ |
| --- | --- | --- | --- | --- | --- |
| Li and Stephan (2006) | 158,000 | 8,603,000 | $N_{\text{EUR},b} = 2,200$ for $t_b = 3,400$ generations, then $N_{\text{EUR}} = 1,075,000$ | 0.3950 | 0.5372 |
| Laurent et al. (2011) | 168,490 | 3,589,770 | Exponential growth from $N_{\text{EUR},t} = 16,550$ to $N_{\text{EUR},0} = 1,224,378$ | 0.5165 | 0.4970 |
| Duchen et al. (2013) | 194,984 | 4,975,360 | Exponential growth from $N_{\text{EUR},t} = 16,982$ to $N_{\text{EUR},0} = 3,122,470$ | 0.4912 | 0.5092 |

For Li and Stephan (2006), values for the model in their Figure 1A appear on pp. 1582-1583; African population size is labeled 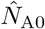 by Li and Stephan (2006) (p. 1582), current European population size is *N*_E0_ (p. 1583), bottleneck European population size is 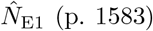, and time variables are 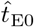 for bottleneck length and 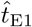 for post-bottleneck interval length (p. 1583). In their inference algorithm, although the data are from the X chromosome, the authors use a coalescent process with *N* diploid individuals and a parametrization Θ = 4*Nµ*. In our calculations in Section 3, we have also assumed that the coalescent process has *N* diploid individuals and 2*N* lineages. Because the parameterization of Li and Stephan (2006) matches our parameterization in Section 3, we use the values from Li and Stephan (2006) directly.

For Duchen et al. (2013), we extracted parameter values for their model C (defined in their Table S2) for the African population from their Table 4 (*N*_Ac_) and for the European population from their Table 5 (*N*_Ea_ and *N*_Ec_ immediately after the split and at the present time, respectively, and time *T*_AE_ since the split), exponentiating values reported logarithmically. The tables of Duchen et al. (2013) report numbers of diploid individuals in a population; hence, we use their population size values directly to calculate 𝔼[Ψ] and 𝕍[Ψ].

For Laurent et al. (2011), we extracted estimates of parameter values from the X-chromosome column in their Table 3 for the model in their Figure 1. Laurent et al. (2011) assumed equal proportions of males and females in the population, explicitly considering the X chromosome, so that the total number of lineages in a population of size *N* is 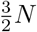, and Θ = 3*Nµ*. To match the diploid autosomal parameterizations under which we derived our theoretical expressions in Section 3, we re-scaled population sizes reported in Table 3 of Laurent et al. (2011) by multiplying them by 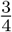, and we then used them to calculate 𝔼[Ψ] and 𝕍[Ψ]; this procedure is equivalent to rederiving our expressions in Section 3 with 3*N* in place of 4*N* and then inserting the *N* values from Laurent et al. (2011) directly.

For the demographic parameters of Li and Stephan (2006) that used a simple bottleneck model, we used eq. (30) with *N*_A_ = *N*_EUR_, *N*_b_ = *N*_EUR,b_, *N*_B_ = *N*_AFR_, and *t*_b_ = *t*_b_ from Table 1. For the demographic parameters of Laurent et al. (2011) and Duchen et al. (2013), which used an exponential growth model, we used eq. (28) with *N*_A,0_ = *N*_EUR,0_, *N*_A,*t*_ = *N*_EUR,*t*_, *N*_B_ = *N*_AFR_, and the exponential growth rate *r*_A_ computed from European population sizes using eq. (23).

The values of 𝔼[Ψ] and 𝕍[Ψ] computed for each set of parameters are shown in Table 1. For the Laurent et al. (2011) and Duchen et al. (2013) demographies, the value we expect from coalescent theory—𝔼[Ψ] in Table 1—lies inside the 95% bootstrap interval for the value obtained directly from data in eq. (37).

The variance in our empirical calculation (eq. (38)) closely matches the values of 𝕍[Ψ] implied in Table 1 by the demographic models, with all three values lying in the 95% bootstrap interval.

## 7 Discussion

### 7.1 Summary

We have examined the directionality index Ψ as a random variable under coalescent models of two populations with a shared demographic history (Section 2). Using this formulation, we have derived exact values for the expectation 𝔼[Ψ] and variance 𝕍[Ψ] of the directionality index for four parameterizations of a population split demography (Section 3). We have explored the behavior of the expectation and variance, showing the dependence of Ψ on demographic parameters and identifying parameter regions for which a positive value of *ψ* does not necessarily mean that the “A” population is more distant from the source of a range expansion (Section 4). Our expression for 𝕍[Ψ] also allowed us to connect the sample variance of *ψ* across many sites to the size of the ancestral population (Section 5). Finally, in Section 6, we showed how our theoretical results can be used to compare the predictions of demographic models with empirical observations.

Our explorations of the theoretical behavior of *ψ* in Section 4 show that in a sample of size 4 lineages, 𝔼[Ψ] tends to be more sensitive to changes in the bottleneck strength *s* and derived population sizes *N*_A_ and *N*_B_ than to the time *t* since the population split (Figure 4a-c). The variance 𝕍[Ψ], however, increases quickly with increasing *t* (Figure 4d-f). These results are informative for considering the effects of bottlenecks on empirical values of *ψ*. For example, in a model in which a bottleneck is ancient, the variance of Ψ would be larger compared to a model with a recent bottleneck.

Expressions for 𝔼[Ψ] and 𝕍[Ψ] (eqs. (20), (26), (30) and (32)) do not depend on *N*_C_, the ancestral population size. This insight suggests that predictions about range expansions under the model are largely unaffected by events in the shared history of the two populations. However, *N*_C_ does affect the number of SNPs available for empirical calculation. When data from many sites are used to compute *ψ*, the expected number of shared SNPs in the calculation is proportional to Θ_C_ = 4*N*_C_*µ*; for small *N*_C_, we might not observe enough shared SNPs for the empirical computation of *ψ* to accurately reflect a model-based prediction.

### 7.2 Comparison of theoretical and empirical *ψ* for *D. melanogaster*

In an application to data from *Drosophila melanogaster*, empirical evaluation of *ψ* revealed a positive value in a scenario with a European population in the role of the “A” population and an African population in the role of the “B” population. This observation is consistent with the higher level of drift in European populations of *D. melanogaster* than in African populations: the analysis accords with the general understanding of *D. melanogaster* demographic history. Further, for two of three *D. melanogaster* modeling studies, the empirical value of *ψ* matched the predictions for 𝔼[Ψ] from the demographic models (Table 1).

### 7.3 Connections

Our results recapitulate some of the insights of the branching process analysis of the discrete-time expansion model of Peter and Slatkin (2015). In that model, the expectation of Ψ was found to be 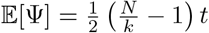 where *N* is the population size of each deme, *k* is the founder population size during settlement of a new deme, and *t* is the settlement time of the *t*th deme, the integer time variable that counts sequential founder events from the origin to the most recently settled deme. This expression shows that 𝔼[Ψ] increases with smaller values for sizes of the founder populations (*k*) and with the number of founder events (*t*).

In our formulations, the founder population size corresponds to the initial population size after the split *N*_A,*t*_ in the exponential growth model (Section 3.2), the bottleneck population size *N*_b_ in the bottleneck model (Section 3.3), and the reciprocal of the bottleneck strength *s* in the instantaneous bottleneck model (Section 3.4). In these cases, we have observed a similar pattern in the magnitude of 𝔼[Ψ], which increases with smaller *N*_A,*t*_ (eq. (26a)), smaller *N*_b_ (eq. (30a)), or larger *s* (eq. (32a), Figure 4).

The role of the time variable *t*, however, differs between the model of Peter and Slatkin (2015) and our analysis. In particular, the linear chain of many populations in Peter and Slatkin (2015) gives rise to a linear dependence of 𝔼[Ψ] on *t*, whereas our two-population model produces a nonlinear dependence of 𝔼[Ψ] on *t* due to interactions among various demographic parameters (Figure 6).

Our study follows a similar spirit to the work of DeGiorgio et al. (2011), who derived expressions for the distribution of pairwise coalescence times in a serial founder model with a sequence of multiple bottlenecks. Many population-genetic statistics are functions of expected pairwise coalescence times, among them *F*_*ST*_ (Slatkin, 1991) and *f*_4_ (Peter, 2016). Because our study uses ratios of certain expected branch lengths rather than the pairwise coalescence times themselves, our expressions for 𝔼[Ψ] and 𝕍[Ψ] are perhaps more closely connected to analyses that focus on internal and external branch length computations and other coalescent-based branch length ratios (Fu & Li, 1993; Ferretti et al., 2017; Alimpiev & Rosenberg, 2022).

An additional connection to other coalescent studies (Tajima, 1983; Takahata & Nei, 1985; Takahata & Slatkin, 1990; Szpiech & Rosenberg, 2011; Rosenberg, 2013; Guerra & Nielsen, 2022; Peter, 2022) is that we have focused on the case of 4 sampled lineages. In many problems, the 4-lineage analysis is the simplest non-trivial case, it can be studied analytically, and it provides insights useful for larger samples.

### 7.4 Further work

Our models are focused on pairs of populations, bottlenecks, and infinitely-many-sites mutation. Extended models could potentially consider additional phenomena; for example, recurrent and reverse mutation, the influence of natural selection on coalescence times for some sites, linkage among sites, and demographies that allow migration after population divergence. In the case of migration, a more recent study of than those that underlie Table 1 suggests a high rate of back-migration of *D. melanogaster* from Europe to Africa (Arguello et al., 2019), so that predictions for *ψ* in models that include migration would be meaningful. One approach to considering migration with *ψ* is an extension of the discrete-time expansion model of Peter and Slatkin (2015). Including migration between demes after founding events and exploring its impact on *ψ* is possible in a simulation-based extension of their model (Kemppainen et al., 2024).

In the framework of our theoretical analysis with four lineages and two populations, migration would allow for private mutations that appeared in population A to be introduced into population B, decreasing the value of *ψ*. Topologies that are more likely to generate shared mutations (such as *δ, ε, ζ* in Figure 2) would be observed more often, due to lineages being transported between populations in the time since the split between populations A and B, altering the values of 𝔼[Ψ] and 𝕍[Ψ]. The theory could potentially be pursued by adding *ψ* to coalescent models that allow for post-divergence migration (Wakeley, 1996; Teshima & Tajima, 2002; Rosenberg & Feldman, 2002; Wilkinson-Herbots, 2008, 2012; Hobolth et al., 2011).

## Data and code

The genetic sequences for *Drosophila melanogaster* X-chromosomal loci were obtained from the European Nucleotide Archive (ebi.ac.uk/ena), sequence IDs AJ568984 to AJ571588, and AJ568984 to AJ571588. Code to perform the computation of *ψ* from data and generate figures and tables in this paper is available at github.com/EgorLappo/coalescent-psi.

## Acknowledgments

We thank Ben Peter for help with a key aspect of the calculation.

## References

Achaz, G. (2009). Frequency spectrum neutrality tests: One for all and all for one. Genetics, 183(1), 249–258. doi: 10.1534/genetics.109.104042.

Alimpiev, E., & Rosenberg, N. A. (2022). A compendium of covariances and correlation coefficients of coalescent tree properties. Theoretical Population Biology, 143, 1–13. doi: 10.1016/j.tpb.2021.09.008.

Arguello, J. R., Laurent, S., & Clark, A. G. (2019). Demographic history of the human commensal Drosophila melanogaster. Genome Biology and Evolution, 11(3), 844–854. doi: 10.1093/gbe/evz022.

Bunnefeld, L., Frantz, L. A. F., & Lohse, K. (2015). Inferring bottlenecks from genome-wide samples of short sequence blocks. Genetics, 201(3), 1157–1169. doi: 10.1534/genetics.115.179861.

Caicedo, A. L., Williamson, S. H., Hernandez, R. D., Boyko, A., Fledel-Alon, A., York, T. L., Polato, N. R., Olsen, K. M., Nielsen, R., McCouch, S. R., Bustamante, C. D., & Purugganan, M. D. (2007). Genome-wide patterns of nucleotide polymorphism in domesticated rice. PLoS Genetics, 3(9), e163. doi: 10.1371/journal.pgen.0030163.

DeGiorgio, M., Degnan, J. H., & Rosenberg, N. A. (2011). Coalescence-time distributions in a serial founder model of human evolutionary history. Genetics, 189(2), 579–593. doi: 10.1534/genetics.111.129296.

DeGiorgio, M., Jakobsson, M., & Rosenberg, N. A. (2009). Explaining worldwide patterns of human genetic variation using a coalescent-based serial founder model of migration outward from Africa. Proceedings of the National Academy of Sciences, 106(38), 16057–16062. doi: 10.1073/pnas.0903341106.

Deshpande, O., Batzoglou, S., Feldman, M. W., & Cavalli-Sforza, L. L. (2009). A serial founder effect model for human settlement out of Africa. Proceedings of the Royal Society B: Biological Sciences, 276(1655), 291–300. doi: 10.1098/rspb.2008.0750.

Duchen, P., Živković, D., Hutter, S., Stephan, W., & Laurent, S. (2013). Demographic inference reveals African and European admixture in the North American Drosophila melanogaster population. Genetics, 193(1), 291–301. doi: 10.1534/genetics.112.145912.

Durrett, R. (2008). Probability models for DNA sequence evolution (2., Ed). xsSpringer.

Edgar, R. C. (2004). MUSCLE: Multiple sequence alignment with high accuracy and high throughput. Nucleic Acids Research, 32(5), 1792–1797. doi: 10.1093/nar/gkh340.

Edmonds, C. A., Lillie, A. S., & Cavalli-Sforza, L. L. (2004). Mutations arising in the wave front of an expanding population. Proceedings of the National Academy of Sciences, 101(4), 975–979. doi: 10.1073/pnas.0308064100.

Excoffier, L., Dupanloup, I., Huerta-Sánchez, E., Sousa, V. C., & Foll, M. (2013). Robust demographic inference from genomic and SNP data. PLoS Genetics, 9(10), e1003905. doi: 10.1371/journal.pgen.1003905.

Excoffier, L., Foll, M., & Petit, R. J. (2009). Genetic consequences of range expansions. Annual Review of Ecology, Evolution, and Systematics, 40(1), 481–501. doi: 10.1146/annurev.ecolsys.39.110707.173414.

Excoffier, L., & Ray, N. (2008). Surfing during population expansions promotes genetic revolutions and structuration. Trends in Ecology & Evolution, 23(7), 347–351. doi: 10.1016/j.tree.2008.04.004.

Ferretti, L., Ledda, A., Wiehe, T., Achaz, G., & Ramos-Onsins, S. E. (2017). Decomposing the site frequency spectrum: The impact of tree topology on neutrality tests. Genetics, 207 (1), 229–240. doi: 10.1534/genetics.116.188763.

Ferretti, L., Perez-Enciso, M., & Ramos-Onsins, S. (2010). Optimal neutrality tests based on the frequency spectrum. Genetics, 186(1), 353–365. doi: 10.1534/genetics.110.118570.

Fu, Y. X., & Li, W. H. (1993). Statistical tests of neutrality of mutations. Genetics, 133(3), 693–709. doi: 10.1093/genetics/133.3.693.

Galtier, N., Depaulis, F., & Barton, N. H. (2000). Detecting bottlenecks and selective sweeps from DNA sequence polymorphism. Genetics, 155(2), 981–987. doi: 10.1093/genetics/155.2.981.

Glinka, S., Ometto, L., Mousset, S., Stephan, W., & De Lorenzo, D. (2003). Demography and natural selection have shaped genetic variation in Drosophila melanogaster: A multi-locus approach. Genetics, 165(3), 1269–1278. doi: 10.1093/genetics/165.3.1269.

Guerra, G., & Nielsen, R. (2022). Covariance of pairwise differences on a multi-species coalescent tree and implications for F_ST_. Philosophical Transactions of the Royal Society B: Biological Sciences, 377 (1852), 20200415. doi: 10.1098/rstb.2020.0415.

Gutenkunst, R. N., Hernandez, R. D., Williamson, S. H., & Bustamante, C. D. (2009). Inferring the joint demographic history of multiple populations from multidimensional SNP frequency data. PLoS Genetics, 5(10), e1000695. doi: 10.1371/journal.pgen.1000695.

Hallatschek, O., & Nelson, D. R. (2008). Gene surfing in expanding populations. Theoretical Population Biology, 73(1), 158–170. doi: 10.1016/j.tpb.2007.08.008.

Hobolth, A., Andersen, L. N., & Mailund, T. (2011). On computing the coalescence time density in an isolation-with-migration model with few samples. Genetics, 187 (4), 1241–1243. doi: 10.1534/genetics.110.124164.

Ioannidis, A. G., Blanco-Portillo, J., Sandoval, K., Hagelberg, E., Barberena-Jonas, C., Hill, A. V. S., Rodríguez-Rodríguez, J. E., Fox, K., Robson, K., Haoa-Cardinali, S., Quinto-Cortés, C. D., Miquel-Poblete, J. F., Auckland, K., Parks, T., Sofro, A. S. M., Ávila-Arcos, M. C., Sockell, A., Homburger, J. R., Eng, C., … Moreno-Estrada, A. (2021). Paths and timings of the peopling of Polynesia inferred from genomic networks. Nature, 597 (7877), 522–526. doi: 10.1038/s41586-021-03902-8.

Kemppainen, P., Schembri, R., & Momigliano, P. (2024). Boundary effects cause false signals of range expansions in population genomic data. Molecular Biology and Evolution, 41(5), msae091. doi: 10.1093/molbev/msae091.

Klopfstein, S., Currat, M., & Excoffier, L. (2006). The fate of mutations surfing on the wave of a range expansion. Molecular Biology and Evolution, 23(3), 482–490. doi: 10.1093/molbev/msj057.

Laurent, S. J. Y., Werzner, A., Excoffier, L., & Stephan, W. (2011). Approximate bayesian analysis of Drosophila melanogaster polymorphism data reveals a recent colonization of southeast Asia. Molecular Biology and Evolution, 28(7), 2041–2051. doi: 10.1093/molbev/msr031.

Li, H., & Stephan, W. (2006). Inferring the demographic history and rate of adaptive substitution in Drosophila. PLoS Genetics, 2(10), e166. doi: 10.1371/journal.pgen.0020166.

Liu, X., & Fu, Y.-X. (2020). Stairway plot 2: Demographic history inference with folded SNP frequency spectra. Genome Biology, 21(1), 280. doi: 10.1186/s13059-020-02196-9.

Marchi, N., Schlichta, F., & Excoffier, L. (2021). Demographic inference. Current Biology, 31(6), R276–R279. doi: 10.1016/j.cub.2021.01.053.

Nielsen, R., Hubisz, M. J., Hellmann, I., Torgerson, D., Andrés, A. M., Albrechtsen, A., Gutenkunst, R., Adams, M. D., Cargill, M., Boyko, A., Indap, A., Bustamante, C. D., & Clark, A. G. (2009). Darwinian and demographic forces affecting human protein coding genes. Genome Research, 19(5), 838–849. doi: 10.1101/gr.088336.108.

Nielsen, R., Williamson, S., Kim, Y., Hubisz, M. J., Clark, A. G., & Bustamante, C. (2005). Genomic scans for selective sweeps using SNP data. Genome Research, 15(11), 1566–1575. doi: 10.1101/gr.4252305.

Ometto, L., Glinka, S., De Lorenzo, D., & Stephan, W. (2005). Inferring the effects of demography and selection on Drosophila melanogaster populations from a chromosome-wide scan of DNA variation. Molecular Biology and Evolution, 22(10), 2119–2130. doi: 10.1093/molbev/msi207.

Peischl, S., & Excoffier, L. (2016). Expansion load: Recessive mutations and the role of standing genetic variation. In Invasion genetics (pp. 218–231). Wiley. doi: 10.1002/9781119072799.ch13.

Peter, B. M. (2016). Admixture, population structure, and F -statistics. Genetics, 202(4), 1485–1501. doi: 10.1534/genetics.115.183913.

Peter, B. M. (2022). A geometric relationship of F _2_, F _3_ and F _4_ -statistics with principal component analysis. Philosophical Transactions of the Royal Society B: Biological Sciences, 377 (1852), 20200413. doi: 10.1098/rstb.2020.0413.

Peter, B. M., & Slatkin, M. (2013). Detecting range expansions from genetic data. Evolution, 67 (11), 3274–3289. doi: 10.1111/evo.12202.

Peter, B. M., & Slatkin, M. (2015). The effective founder effect in a spatially expanding population. Evolution, 69(3), 721–734. doi: 10.1111/evo.12609.

Pool, J. E., Hellmann, I., Jensen, J. D., & Nielsen, R. (2010). Population genetic inference from genomic sequence variation. Genome Research, 20(3), 291–300. doi: 10.1101/gr.079509.108.

Puckett, E. E., & Munshi-South, J. (2019). Brown rat demography reveals pre-commensal structure in eastern Asia before expansion into southeast Asia. Genome Research, 29(5), 762–770. doi: 10.1101/gr.235754.118.

Ramachandran, S., Deshpande, O., Roseman, C. C., Rosenberg, N. A., Feldman, M. W., & Cavalli-Sforza, L. L. (2005). Support from the relationship of genetic and geographic distance in human populations for a serial founder effect originating in Africa. Proceedings of the National Academy of Sciences, 102(44), 15942–15947. doi: 10.1073/pnas.0507611102.

Ronen, R., Udpa, N., Halperin, E., & Bafna, V. (2013). Learning natural selection from the site frequency spectrum. Genetics, 195(1), 181–193. doi: 10.1534/genetics.113.152587.

Rosenberg, N. A. (2013). Discordance of species trees with their most likely gene trees: A unifying principle. Molecular Biology and Evolution, 30(12), 2709–2713. doi: 10.1093/molbev/mst160.

Rosenberg, N. A., & Feldman, M. W. (2002). The relationship between coalescence times and population divergence times. In Modern developments in theoretical population genetics (pp. 130–164). Oxford University Press. doi: 10.1093/oso/9780198599623.003.0009.

Schlichta, F., Moinet, A., Peischl, S., & Excoffier, L. (2022). The impact of genetic surfing on neutral genomic diversity. Molecular Biology and Evolution, 39(11), msac249. doi: 10.1093/molbev/msac249.

Slatkin, M., & Hudson, R. R. (1991). Pairwise comparisons of mitochondrial DNA sequences in stable and exponentially growing populations. Genetics, 129(2), 555–562. doi: 10.1093/genetics/129.2.555.

Slatkin, M. (1991). Inbreeding coefficients and coalescence times. Genetical Research, 58(2), 167–175. doi: 10.1017/S0016672300029827.

Slatkin, M., & Excoffier, L. (2012). Serial founder effects during range expansion: A spatial analog of genetic drift. Genetics, 191(1), 171–181. doi: 10.1534/genetics.112.139022.

Stephan, W., & Li, H. (2007). The recent demographic and adaptive history of Drosophila melanogaster. Heredity, 98(2), 65–68. doi: 10.1038/sj.hdy.6800901.

Szpiech, Z. A., & Rosenberg, N. A. (2011). On the size distribution of private microsatellite alleles. Theoretical Population Biology, 80(2), 100–113. doi: 10.1016/j.tpb.2011.03.006.

Tajima, F. (1983). Evolutionary relationship of DNA sequences in finite populations. Genetics, 105(2), 437–460. doi: 10.1093/genetics/105.2.437.

Takahata, N., & Nei, M. (1985). Gene genealogy and variance of interpopulational nucleotide differences. Genetics, 110(2), 325–344. doi: 10.1093/genetics/110.2.325.

Takahata, N., & Slatkin, M. (1990). Genealogy of neutral genes in two partially isolated populations. Theoretical Population Biology, 38(3), 331–350. doi: 10.1016/0040-5809(90)90018-Q.

Teshima, K. M., & Tajima, F. (2002). The effect of migration during the divergence. Theoretical Population Biology, 62(1), 81–95. doi: 10.1006/tpbi.2002.1580.

Thornton, K., & Andolfatto, P. (2006). Approximate bayesian inference reveals evidence for a recent, severe bottleneck in a Netherlands population of Drosophila melanogaster. Genetics, 172(3), 1607–1619. doi: 10.1534/genetics.105.048223.

Wakeley, J. (1996). Pairwise differences under a general model of population subdivision. Journal of Genetics, 75(1), 81–89. doi: 10.1007/BF02931753.

Wakeley, J., & Hey, J. (1997). Estimating ancestral population parameters. Genetics, 145(3), 847–855. doi: 10.1093/genetics/145.3.847.

Wilkinson-Herbots, H. M. (2008). The distribution of the coalescence time and the number of pairwise nucleotide differences in the “isolation with migration” model. Theoretical Population Biology, 73(2), 277–288. doi: 10.1016/j.tpb.2007.11.001.

Wilkinson-Herbots, H. M. (2012). The distribution of the coalescence time and the number of pairwise nucleotide differences in a model of population divergence or speciation with an initial period of gene flow. Theoretical Population Biology, 82(2), 92–108. doi: 10.1016/j.tpb.2012.05.003.

Zhan, S., Zhang, W., Niitepõld, K., Hsu, J., Haeger, J. F., Zalucki, M. P., Altizer, S., De Roode, J. C., Reppert, S. M., & Kronforst, M. R. (2014). The genetics of monarch butterfly migration and warning colouration. Nature, 514(7522), 317–321. doi: 10.1038/nature13812.

